# Niche complementarity among pollinators increases community-level plant reproductive success

**DOI:** 10.1101/629931

**Authors:** Ainhoa Magrach, Francisco P. Molina, Ignasi Bartomeus

**Affiliations:** Basque Centre for Climate Change-BC3, Edif. Sede 1, 1º, Parque Científico UPV-EHU, Barrio Sarriena s/n, 48940, Leioa, Spain; IKERBASQUE, Basque Foundation for Science, María Díaz de Haro 3, 48013, Bilbao, Spain; Estación Biológica de Doñana (EBD-CSIC), Avda. Américo Vespucio 26, Isla de la Cartuja, 41092, Sevilla, Spain

**Keywords:** nestedness, niche complementarity, fruit set, pollination, plant-pollinator interactions

## Abstract

Our understanding of how the structure of species interactions shapes natural communities has increased, particularly regarding plant-pollinator interactions. However, research linking pollinator diversity to reproductive success has focused on pairwise plant-pollinator interactions, largely overlooking community-level dynamics. Here, we present one of the first empirical studies linking pollinator visitation to plant reproduction from a community-wide perspective. We use a well-replicated dataset encompassing 16 plant-pollinator networks and data on reproductive success for 19 plant species from Mediterranean shrub ecosystems. We find that statistical models including simple visitation metrics are sufficient to explain the variability observed. However, a mechanistic understanding of how pollinator diversity affects reproductive success requires additional information on network structure. Specifically, we find positive effects of increasing complementarity in the plant species visited by different pollinators on plant reproductive success. Hence, maintaining communities with a diversity of species but also of functions is paramount to preserving plant diversity.

## Introduction

Pollinators provide key services to plants by facilitating pollen flow (Garibaldi *et al* 2013). Declining trends for some pollinator species in some regions (Potts *et al*. 2010, Bartomeus *et al*. 2019) have led researchers to focus on the functional impacts of these changes in pollinator diversity, especially for plant reproductive success (Biesmeijer *et al*. 2006).

Many studies have evaluated reproductive success on individual plant species (Albrecht *et al*. 2012; Thomson 2019), and used relatively simple visitation metrics (e.g., the number of pollinator species visiting a plant or the number of visits they perform) to explain the differences observed (e.g., Bommarco et al 2012). Contrastingly, community-level analyses remain scarce (Bennett *et al*. 2018). Yet plants and pollinators do not interact in isolation but are embedded within larger networks of interactions encompassing other plant and pollinator species (Memmot *et al* 2004). We are thus missing an important part of the picture, including direct interactions between the whole ensemble of plants and pollinators, but also indirect ones between species within one guild (e.g., plants) through their shared resources (Pauw 2013; Carvalheiro *et al*. 2014; Lázaro *et al*. 2014; Mayfield & Stouffer 2017; Johnson & Bronstein 2019). Understanding how changes in pollinator diversity and community structure affect ecosystem functioning is thus a major challenge that requires attention.

The few studies that have analyzed the effects of pollinator diversity on reproductive success at the community level have mainly used experimental setups. As an example, a study that experimentally recreated a plant community with 9 plant species and differing levels of pollinator diversity, found a positive effect of pollinator species diversity on seed set, but also an important effect of niche complementarity between pollinators, a measure of community structure (Fründ *et al*. 2013). These findings show that not only the diversity of species present, but also the diversity of roles they play and thus the way in which a community is structured are determinant factors of ecosystem functions.

Indeed, theoretical research has long suggested that the structure of multitrophic communities has an effect for ecosystem functioning (reviewed in Thompson *et al*. 2012). This line of research, rooted in niche theory and revamped by food-web studies (MacArthur & Levins 1967; May & MacArthur 1972, Tilman 1982, Godoy *et al* 2018), has greatly advanced theory, but the relationship between structure and function has seldom been tested using empirical data (but see Poisot *et al*. 2013, Kaiser-Bunbury *et al* 2017, Lazaro *et al* 2019). Specifically, a major knowledge gap resides in understanding which aspects of structure determine which aspects of function (Thompson *et al*. 2012). This is because although a network perspective has promised to encapsulate complex ecological mechanisms occurring at the community level – such as indirect interactions (Holt 1977, Abrams *et al* 1998) or niche overlap (Woodward & Hildrew 2002)- less attention has been given to the ways in which these mechanisms relate to observed ecosystem processes (Blüthgen 2010). We are now at a point where we understand some of the emergent patterns characterizing mutualistic interaction networks at the community level, especially in the case of pollination (Bascompte & Jordano 2007). Amongst them is the prevalence of nested structures, i.e., arrangements where specialist species interact with a subset of the species that generalists interact with (Bascompte *et al* 2003). Further, plant-pollinator interaction networks seem to exhibit a relatively high extent of complementary specialization at the community scale, which may be directly related to key ecosystem functions (Blüthgen & Klein 2011). However, the mechanisms by which these attributes affect plant reproduction remain to be understood (Winfree 2013). The time is thus ripe to explore the relationship between community structure and ecosystem functioning empirically, with special emphasis on the underlying ecological mechanisms that drive these relationships.

Here, we present an empirical study linking pollinator visitation and plant reproductive success at the community level. We use a well-replicated dataset encompassing plant-pollinator interaction networks collected at 16 sites coupled with data on the reproductive success of 19 plant species recorded in Mediterranean shrub ecosystems. Our study focuses on understanding whether adding information on selected interaction network structure indices to previously used simple visitation metrics (e.g., the number and diversity of pollinator species visiting a plant species) aids in better explaining the differences observed in communitywide reproductive success. In doing so, we conducted our analyses focusing on reproductive success at two different levels: (i) at the species level by considering the association between the position of a focal species within the larger network and its link to individual reproductive success, and (ii) at the site level, by evaluating how attributes that describe the whole site might affect average values of reproductive success for all species measured within one particular site. Specifically, our study focuses on how the interplay between the complementarity in plant species visited by different pollinators, and the redundancy in this function relate to reproductive success. Plant reproductive success requires the delivery of conspecific pollen and thus of a certain degree of niche complementarity (Blüthgen & Klein 2011). Yet, greater values of redundancy in species functions (e.g., that provided by nested structures), are thought to promote species diversity (Bastolla *et al*. 2009) and stability (Thébault & Fontaine 2010) within plant-pollinator networks. At present, we do not know how either of these network characteristics affects the functions performed by pollinators.

Our results suggest that models including information on simple visitation metrics alone are able to explain differences in reproductive success. However, a mechanistic understanding requires additional information on network structure, notably information on the complementarity between the niches occupied by different pollinator species. Specifically, we find a positive effect of increasing niche complementarity between pollinators on plant reproductive success.

## Methods

### Plant pollinator interactions

Our study was conducted in SW Spain within the area of influence of Doñana National Park (Fig. S1). Sites were located within similar elevations (ranging from 50 to 150 m a.s.l.), similar habitat and soil types, and presented similar plant composition (plant mean Sørensen beta-diversity among sites = 0.41), reducing potential confounding factors. We surveyed 16 Mediterranean woodland patches with an average distance of 7 km between them (min= 3 km, max= 46.5 km). Each site was surveyed every two weeks for a total of 7 times during the flowering season of 2015 (from February to May) following a 100-m x 2 m transect for 30 mins. Along each transect, we identified all plant species and recorded all the floral visitors that landed on their flowers and touched the plant’s reproductive parts. Only floral visitors (from now on referred to as pollinators) that could not be identified in the field were captured, stored and identified in the laboratory by FPM and another expert entomologist (see acknowledgements). All surveys were done under similar weather conditions, avoiding windy or rainy days, during mornings and afternoons with the sampling order being established randomly. Within each transect every 10 m we surveyed a 2×2 m quadrant where the number of flowers per species were counted, i.e., 10 quadrats per transect which makes 40m^2^ of area surveyed overall.

### Plant reproductive success

Within each site, we marked 3-12 individuals (mean ± SD: 6.49 **±** 2.37) belonging to 1-6 plant species (mean ± SD: 4.06 **±** 1.69, Table S2). For each individual, at the end of the season, we recorded fruit set (i.e. the proportion of flowers that set fruit), the average number of seeds per fruit and the average fruit and seed weight per fruit (1-36 fruits subsampled; mean ± SD: 11.17 **±** 6.85, Table S3). These last two variables show a strong correlation (Pearson correlation= 0.89), and thus we only present results on fruit weight. Our survey included a total of 19 different totally or partially self-incompatible plant species that depend on pollinators to maximize their reproduction (Table S4) across our 16 sites. All plant species were common and widespread shrubs. Individuals were selected depending on the presence of flowers during the sampling events. We also calculated the average reproductive success at the site level by averaging values of reproductive success obtained for each species.

### Data analyses

To evaluate the sampling completeness, we estimated the asymptotic number of species of plants, pollinators and interactions present (Chao *et al*. 2009), a non-parametric estimator of species richness for abundance data. This estimator includes non-detected species and allowed us to calculate the proportion detected with our original data. We used Chao 1 asymptotic species richness estimators (Chao *et al*. 2009) and estimated the richness of pollinators, plants and plant–pollinator links accumulated as sampling effort increased up to 100% sampling coverage using package iNEXT (Hsieh *et al*. 2016) within the R environment (R Development Core Team 2011). We then extracted the values covered by our sampling.

To evaluate differences in network structure between communities, we constructed plant-pollinator interaction networks by pooling the data for the 7 rounds of sampling. We thus obtained one interaction network per site, representing the number of individuals of different pollinator species recorded visiting each different plant species. For each network, we extracted a series of relevant network metrics at the species and site levels.

Additionally, we checked for spatial autocorrelation in our data using Mantel correlograms. Autocorrelation values were non-significant for all variables, except for pollinator richness where we have a small but significant effect at small spatial scales (Fig. S2). Hence, we treat each site as independent in our analysis.

### Species-level network analysis

At the species level, we focused on attributes defining the position of a focal plant species within the larger community. As such, we considered two metrics providing complementary non-redundant information: (i) average niche overlap in terms of pollinators between a focal plant species and each of the other plant species in the community, and (ii) the contribution to nestedness of each individual plant species. Niche overlap estimates the potential indirect interactions between plant species through shared resources (in this case pollinators) and the potential for increased heterospecific pollen deposition (Arceo-Gómez *et al* 2019). We calculated it as the average overlap in pollinator species visiting a focal plant and each of the other plants in the community using the Morisita overlap index, a measure of similarity between two sets of data (Zhang 2016). A plant species’ contribution to nestedness is calculated by comparing the nestedness observed in a given community to that generated by randomizing the interactions in which a focal species is involved. Species that show important contributions to overall nestedness will have values >0, while species that do not contribute to overall nestedness wil show values <0 (Saavedra *et al* 2011).

### Site-level network analysis

At the site level, we followed the same logic as the one presented at the species level. We also calculated two network metrics providing complementary non-redundant information. In this case, we focused on nestedness, a measure of the redundancy in the plants visited by different pollinators, and pollinator niche complementarity, a measure of the complementarity in plant species visited by different pollinator species.

Nestedness is the property by which specialists interact with a subset of the species that generalists interact with (Bascompte *et al*. 2003). Although there is an ongoing debate in the literature (e.g., James *et al* 2012), some theoretical studies have found that nested networks are more stable and resilient to perturbations because nestedness promotes a greater diversity by minimizing competition among species in a community (Bastolla *et al*. 2009). However, many network attributes vary with network size and complexity (Blüthgen *et al*. 2006). In the case of nestedness, we know it can be affected by network size and connectance (Song *et al*. 2017). An approach that is often used to correct for this are null models, comparing null-model corrected nestedness values across different networks. However, this approach presents the same issues, as z-scores also change with network size and connectance (Song *et al*. 2017). We thus used a normalized value of the widely used nestedness metric NODF based on binary matrices (Almeida-Neto & Ulrich 2011), ***NODF_C_*** (Song *et al*. 2017). This normalized value is calculated as ***N0DF_C_* = *NOD F_n_*/(*C* * *log*(*S*))**, where C is connectance and S is network size, calculated as 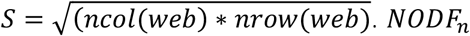 is calculated as ***NODF/max*(*NODF*)**, which is independent of network size and thus comparable across different networks (Song *et al* 2017). To calculate max(NODF) we used a corrected version of the algorithm (Simmons *et al* 2019) whenever possible. Results did not change qualitatively when using the uncorrected version of the algorithm for all sites as both are highly correlated (Spearman correlation = 0.94).

To calculate niche complementarity, we used a community-level measure defined as the total branch length of a dendrogram based on qualitative differences in visitor assemblages between plants (Petchey & Gaston 2007; Devoto *et al*. 2012). All network metrics were calculated using package bipartite (Dormann *et al*. 2009).

### Statistical analyses

To evaluate whether adding information on network structure improves our ability to explain differences in reproductive success - both at the species and the site level - we used generalized linear (GLMs) and generalized linear mixed models (GLMMs) respectively. In both cases we fit three types of models: (i) model 0, a null model with no explanatory variables, (ii) model 1, that only included simple visitation metrics and (iii) model 2 that additionally included information on network structure. These models are meant to be additive, so that the network metrics included are intended to complement rather than substitute the simple metrics traditionally used.

At the species level, response variables included fruit set analyzed using a binomial distribution and the average number of seeds per fruit, and the average fruit weight fitted using normal distributions. The number of seeds per fruit was centered and scaled (i.e., we subtracted column means and divided by standard deviation) to allow meaningful comparisons across species with contrasting life histories. As explanatory variables, model 1 included the number of pollinator species observed, and the visitation rate received by each plant species. Visitation rate was calculated as the total number of visits received by a plant species divided by the average number of flowers of that species found in the 10 2×2 m quadrats per transect. In turn, model 2 added the two network attributes calculated at the species level: average plant niche overlap and contribution to nestedness. For both models, we included plant species identity nested within site and site as random effects to account for multiple individuals of the same plant species measured at each site.

At the site level, response variables were the average reproductive success of all plants surveyed within a site (i.e., average fruit set analyzed using a binomial distribution, average number of seeds per fruit and average fruit weight using a normal distribution). We thus had a single value per site and no random effects are needed. Here, model 1 included total pollinator richness and total pollinator abundance (i.e. number of visits received by all plants within the community) as explanatory variables. Model 2, in turn, added information on network structure by including nestedness and pollinator niche complementarity.

Average values of reproductive success at the site level can be driven by a single plant species. Yet, what will determine the persistence of a diverse plant community, is the presence of some sort of “equity” or evenness in reproductive success across the whole community. We therefore calculated the proportion of species with normalized (between 0 and 1) average fruit set values that were above the 50^th^ percentile as a measure of equity. As any selected threshold is arbitrary, we repeated this using the 25^th^ and 75^th^ percentile thresholds (Byrnes *et al* 2014). We then used the same framework as that used for species and site-level analyses and fit the same models 0, 1 and 2 using equity in reproductive success as response variable and fitting a binomial distribution.

In all cases, we used variance inflation factors to check for collinearity between explanatory variables. Additionally, we ran residual diagnostics to check if model assumptions were met and used the Akaike Information Criterion (AIC) to compare model performance and complexity. Whenever the difference between the AIC of the models was < 2 (***ΔAIC* < 2**), we considered all models equally good (Burnham *et al*. 2011). In the case of mixed models, for comparison, models were fitted by maximum likelihood and then the best model was refitted using restricted maximum likelihood. All predictor variables were standardized prior to analysis. For every model we also calculate the R^2^ value, using the approximation suggested for GLMMs when necessary (Nakagawa *et al* 2017).

Finally, we tested whether the importance of network structure in explaining differences in equity in reproductive success increases with the number of plant species being considered. We expect that when only one plant species is considered the importance of network structure will be negligible, while we expect it to increase as more plant species are considered (up to a maximum number of 6 species which is the maximum we have measured in our study at a particular site).

To test this, we ran a simple simulation in which the number of species considered increased at each step and for each step we re-calculated equity in reproductive success. Instead of drawing plant species randomly for each step, we tested all possible combinations for each plant number level and network, as the number of combinations is small (e.g. for n = 3 plants selected out of 6 there are only 20 possible combinations). Then, we tested if the relationship between equity in reproductive success and niche complementarity (given its importance in determining differences in reproductive success, see Results section) changes as a function of the number of plants considered within our simulated communities. To this end, for each level of species number considered, we randomly selected one of the generated equity values across each of the 16 communities and regressed these 16 values against our network level predictor and extracted the model slope estimates. We repeated this process 1,000 times and averaged all slope estimates. We expect that the more plants considered, the larger the resulting average estimates will be. Note that we only interpret the mean effects, as the variance among different plant number of species considered depends on the initial number of possible combinations.

## Results

Within our sampling we recorded 655 plant-pollinator interactions involving 162 pollinator species and 46 plant species (Table S1). Within the pollinator community the distribution of individuals in different orders was: 92.18% Hymenoptera, 5.69% Diptera, 1.29% Coleoptera and 0.63% Lepidoptera.

Our sampling completeness analyses revealed that our survey was able to capture 17-54% of pollinator species (average = 35%), 43-100% of plant species (average = 80%) and 9-32% of plant-pollinator links (average = 20%; Fig. S3). Our values of sampling completeness were slightly smaller in the case of pollinators, probably as a consequence of the great diversity found in the Mediterranean region and within our study area in particular, a hotspot of insect diversity (Nieto *et al*. 2014).

### Species-level analyses

At the species level, in the case of fruit set, our results showed that model 2 had the best fit to our data (lowest AIC value), and fixed effects explained 9% of the variability observed (conditional R^2^ = 17%). We found a positive relationship between fruit set, pollinator species richness, and a network structure metric, the contribution to nestedness of a focal plant within the overall network (Table 1, Fig. 1, Fig. S4).

**Table 1.**
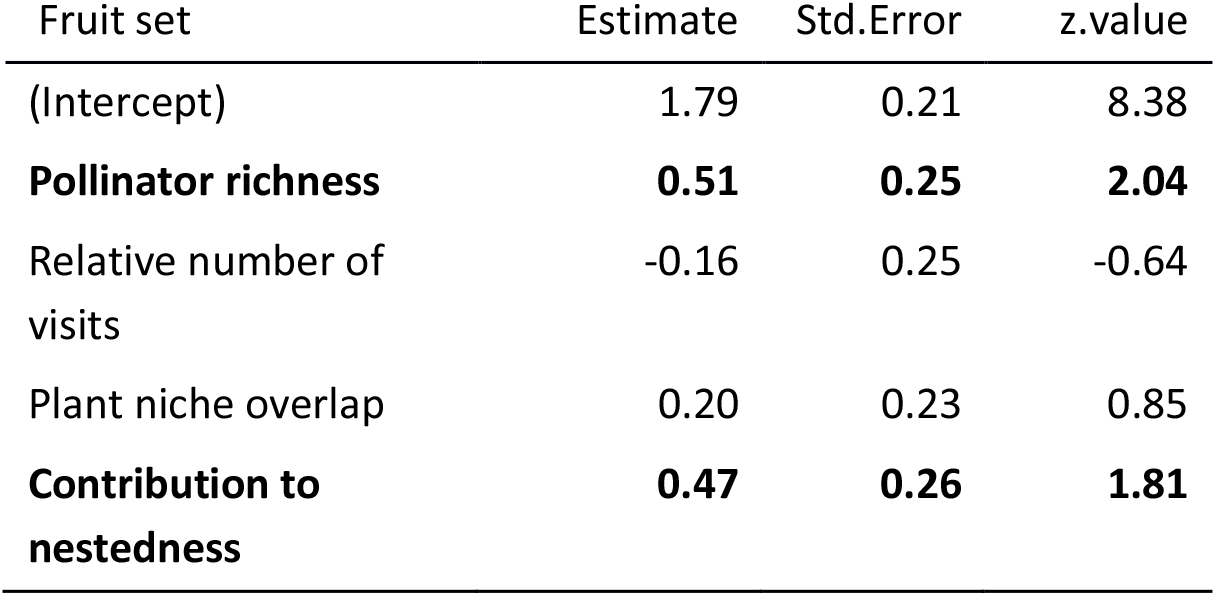
Results of GLMM showing the association between simple visitation and network structure metrics and species-level fruit. Bold letters indicate variables with large effects (see Figure S4 for estimate confidence intervals).

**Figure 1.**
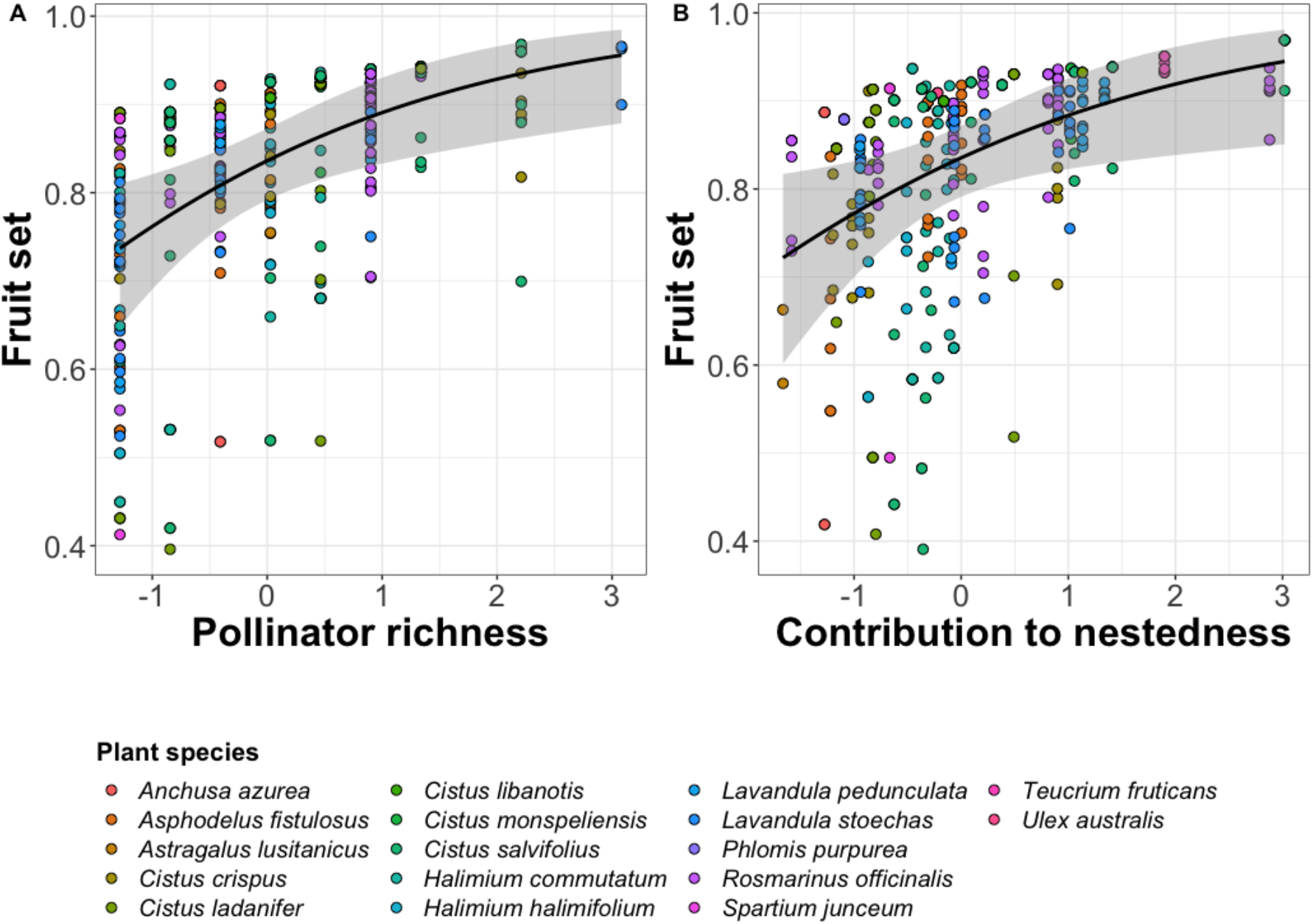
Partial residual plots showing the effect of A) pollinator species richness and B) the contribution to nestedness of each plant species on fruit set. Dots represent each of the individuals sampled for each species within each site.

For the average number of seeds per fruit at the species level as well as for fruit weight, our results showed that none of the models fitted were better than the null model explaining differences across plant species.

### Site-level analyses

At the site level, in the case of fruit set and the number of seeds per fruit, we found that both model 1 and 2 were equally good when penalizing for model complexity (i.e.,***ΔAIC < 2***; Burnham *et al* 2011). This suggests model 2 was a good model despite its added complexity, and actually showed a substantially better predictive ability than model 1 (R^2^ = 0.46 for model 2 versus 0.27 for model 1 in the case of fruit set and R^2^ = 0.49 for model 2 versus 0.35 for model 1 in the case of the number of seeds per fruit) and therefore we will comment results for this model only. Specifically, we found that both fruit set and the number of seeds per fruit were positively related to niche complementarity between pollinators (Table 2, Fig. 2, Fig. S5). Additionally, we found a negative association between site-level pollinator richness and average fruit set (Table 2A, Fig. 2, Fig. S5).

**Table 2.** Results of GLM showing associations between simple visitation and network structure metrics and A) site-level average fruit set and B) site-level average number of seeds per fruit based on best model selected. Bold letters indicate variables with large effects.

| A) Fruit set | Estimate | Std. Error | z value |
| --- | --- | --- | --- |
| (Intercept) | **1.20** | **0.15** | **7.79** |
| **Pollinator richness** | **-0.77** | **0.26** | **-2.91** |
| Relative number of visits | -0.12 | 0.19 | -0.66 |
| Nestedness | 0.02 | 0.16 | 0.12 |
| **Pollinator niche complementarity** | **0.40** | **0.26** | **1.58** |

| B) Seeds per fruit | Estimate | Std. Error | t value |
| --- | --- | --- | --- |
| (Intercept) | 45.37 | 8.84 | 5.13 |
| Pollinator richness | 1.56 | 15.80 | 0.10 |
| Relative number of visits | 4.37 | 10.78 | 0.41 |
| Nestedness | 3.94 | 9.80 | 0.40 |
| **Pollinator niche complementarity** | **26.44** | **15.49** | **1.71** |

**Figure 2.**
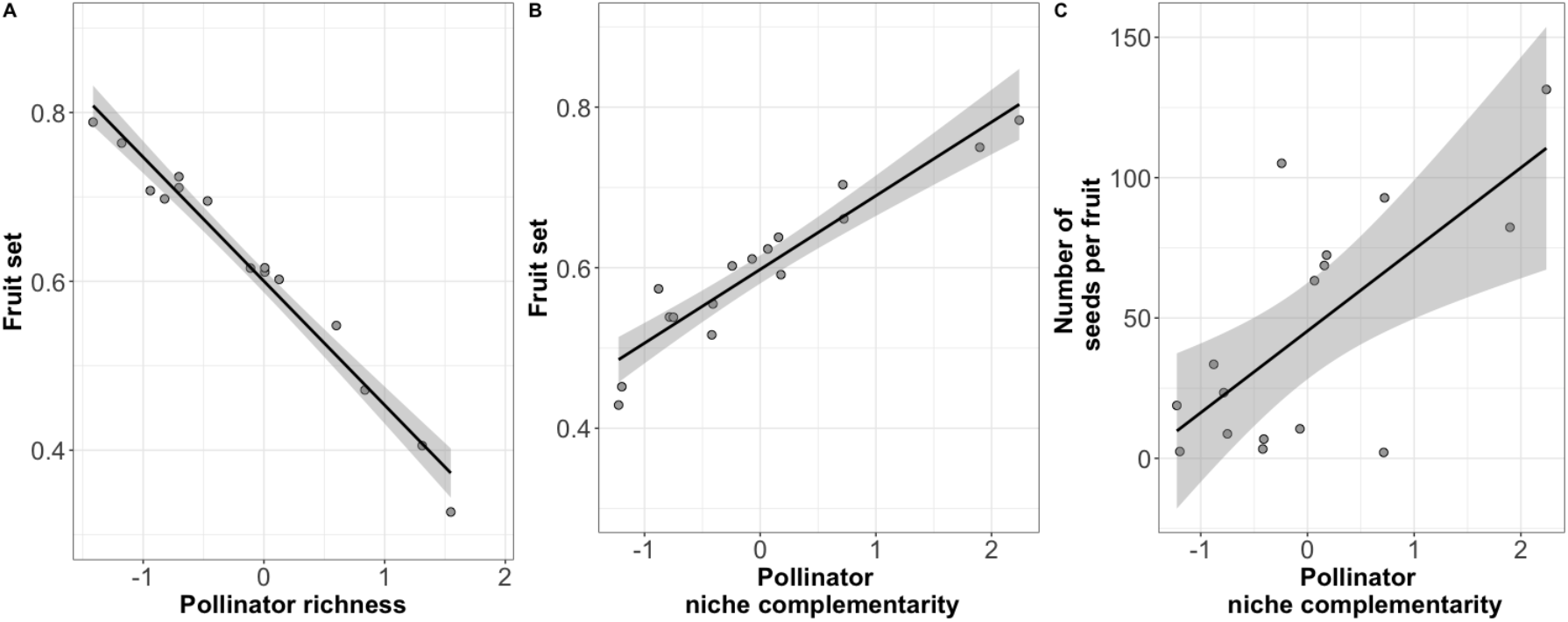
Partial residual plots showing the effect of the single predictor which best explains the variability in site-level reproductive success. A) Shows the effect of pollinator richness, and B) of niche complementarity among pollinator species on site-level average fruit set. C) Shows the effect of niche complementarity among pollinator species on the average number of seeds per fruit at the site level. Dots represent average values of fruit set at the level of the community for all plant species considered (N=16 sites).

In the case of fruit weight, we found that both the null model and model 1 were equally good (i.e.,***ΔAIC < 2***; Burnham *et al* 2011). Model 1, i.e., that only including simple visitation metrics, showed an R^2^ of 0.23. In this case, we found a positive link with site-level pollinator richness (Table S5A, Figs. S5-S6). This association was maintained even after removing a site that has a particularly large pollinator richness value (Table S5B, Fig. S7, Fig. S5).

### Equity in fruitset

When evaluating the relationship between community composition and network structure on equity in reproductive success across the different species within a community, we found that using the 50^th^ percentile all models were equally good (i.e.,***ΔAIC < 2***; Burnham *et al* 2011), but none of the variables considered showed any strong associations (Table S6). In the case of the other two thresholds considered (25^th^ and 75^th^ percentiles) model 0, the null model, was the best model.

Within our simulation evaluating the relationship between niche complementarity and equity in reproductive success at increasing number of plant species considered, we found that the link to complementarity became more important as more species were considered (Fig. 3). This importance seemed to reach a plateau. However, this should be further evaluated, as this was the maximum number of species simultaneously observed in a community for our study, which precludes us from simulating further numbers of species.

**Figure 3.**
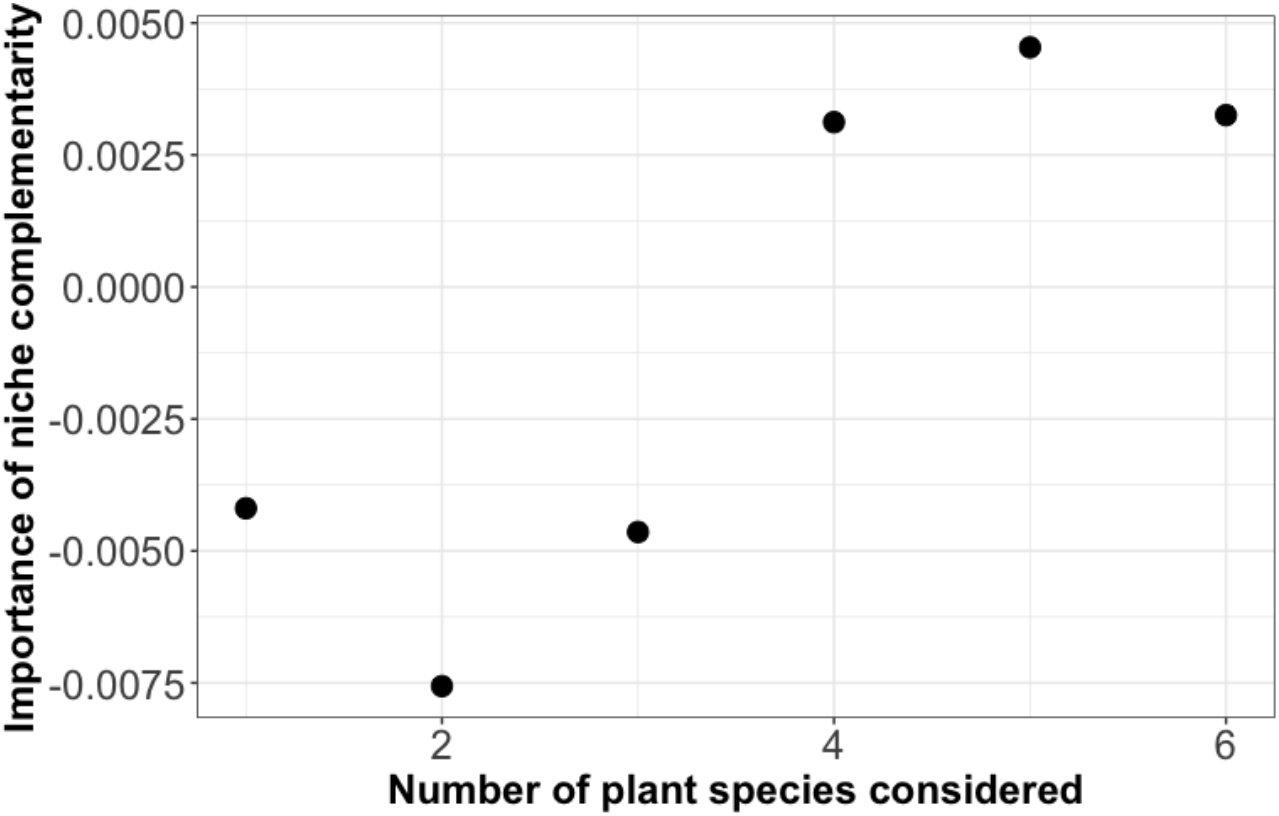
Results of simulation evaluating the importance of niche complementarity in determining differences in equity in reproductive across communities harboring from one to six species. Points represent average values across 1,000 simulated combinations.

## Discussion

The existence of relationships between interaction network structure and ecosystem function have been long hypothesized, yet, the specific mechanisms underlying this relationship remain elusive (Thompson *et al*. 2012). Our results suggest that different aspects of network structure affect different dimensions of ecosystem functioning. Specifically, we find that the contribution to nestedness of a plant species within a community has a positive association with its fruit set. From a plant’s perspective, this indicates that being connected to other plant species via shared pollinators has a positive outcome (e.g. by ensuring a stable pollinator supply through time) rather than a negative one (e.g. via heterospecific pollen transport). At the site level, we find that greater values of niche complementarity between pollinators result in larger average values of reproductive success.

Most of our analyses reveal that model 1 and 2 were equally good, which suggests that the added complexity of measuring the full network of interactions may not pay off for rapid assessments. Hence, simple visitation metrics, such as pollinator richness, might be enough to describe general patterns (Garibaldi *et al*. 2013, 2015). Yet, adding network level information may inform us of the potential ecological mechanisms underlying the processes driving the observed patterns.

Consistent with previous experimental (Fontaine *et al*. 2005; Fründ *et al*. 2013), theoretical (Pauw 2013), and empirical studies (Valdovinos *et al*. 2016, Poisot *et al*. 2013), we find that niche complementarity is key in determining differences in reproductive outputs at the community level, both greater fruit set and larger numbers of seeds per fruit. These results show that reproductive success in plants requires of a certain degree of specialization amongst pollinator species on a particular plant resource in order to avoid the negative effects of inter-specific pollen deposition (e.g., pollen loss, Flanagan *et al*. 2009) or interference with conspecific pollen (Morales & Traveset 2008). However, we also find that some level of redundancy in these functions is needed as revealed by the positive effect of plant niche overlap on the number of seeds per fruit at the species level.

We did not find a relationship between nestedness and any of the reproductive success measures. This metric, widely used across network analysis, does not seem to play a direct role on plant reproductive success. However, our study is limited to a maximum of six common plant species per community, and including more species, especially rare species, might reveal different patterns. Further, although we sampled each site seven times in a randomized order in an attempt to better represent interactions through time, our surveys were able to capture ~20% of interactions given the great diversity of our study system. This could be explaining part of the low effect sizes we find at the species level, where a stronger contribution of pollinator visits is expected given their obligate dependence (Garibaldi *et al* 2013). In addition, it is important to note that plant reproductive success is affected by other environmental variables which we do not attempt to measure in this study and that could explain a large portion of the variability observed.

Our measure of reproductive success at the site level using average values represents an important part of the functions delivered by pollinators to plants. However, average values might mask a great deal of variability amongst plant species, and thus a nuanced view of the effect of pollinators on whole-plant ensembles is needed. This can be captured by the effect of pollinators on equity in reproductive success across plant species. This aspect ensures that reproductive success is equally distributed amongst a larger number of species. Indeed, we know that plant species diversity within a community is largely driven by different types of direct and indirect interactions including those amongst plant species (e.g., resource competition, Goldberg & Barton 1992, or facilitation, Bruno *et al*. 2003), as well as those defining antagonistic (e.g., involving pathogens, Bagchi *et al*. 2010), or mutualistic interactions (e.g, pollinators, Benadi *et al*. 2013; Lanuza *et al*. 2018). However, equitability in reproductive success across species is seldom taken into account, despite increasing theoretical and empirical support to the idea that minimizing fitness differences among species is an important mechanism of species coexistence (Godoy *et al* 2014). In our case, we did not find a strong effect of either simple visitation or network structure metrics on reproductive equity. However, the results of our simulation, shows us that the effect of network structure increases when more than four plant species are considered. This implies that if we were able to measure reproductive success for all the plant species in all the communities (which is not feasible given constraints on sampling effort), we might find that the effects of network structure on equity might be more prevalent.

One of the unexpected results of our analyses is the strong negative relationship between total pollinator richness and fruit set at the site level. One possible explanation for this is that greater richness means greater transfer of heterospecific pollen (Arceo-Gómez *et al* 2019). Another possible explanation to this might be the fact that pollinator richness includes all the pollinators recorded during our sampling efforts, i.e., it includes species that do not pollinate some of the species whose reproductive success was measured. More complex communities with more pollinators, but also with more plant species (Pearson correlation between plant and pollinator richness = 0.42 in our case) may require stabilizing mechanisms that reduce the competition exerted by the dominant plant species. A way to reduce the competition exerted by these dominant species, which are precisely those evaluated in this study, is by reducing their reproductive success (Lanuza *et al* 2018, Stavert *et al* 2019). These ideas open the door to exploring the positive or negative effects of the complete pollinator community on full plant species coexistence, which may be determined by density-dependence effects (Benadi & Pauw 2018). In our case, while fruit set at the site level is negatively related to pollinator richness, it is important to note that fruit set at the species level and fruit weight show the opposite relationship, indicating that this density-dependent effect might only be limiting fruit quantity and not fruit quality. Thus, taking into account the densities of co-flowering plant species may be the next step (Vanbergen *et al*. 2014).

Our study illustrates the challenges of measuring and linking network structure to ecosystem function empirically. There is an ongoing debate as to which network metrics better reflect classic ecological mechanisms, such as niche partitioning or competition (Delmas *et al* 2018). Here, we focus on testing two specific hypotheses, but other structural properties can be explored in the future. Furthermore, the structure of plant-pollinators networks is dynamic due to ecological and evolutionary reasons, but so far, we are only able to characterize it for single snap-shots. Moreover, different aspects of functioning may be important, such as the need to consider the functioning of both trophic levels (Godoy *et al* 2018). In terms of plant reproductive success and the functions performed by pollinators we can measure different aspects, ranging from pollen deposition (the direct pollinator function), to its final effects on plant fitness. Here, we focus on an intermediate stage including fruit quantity and quality, which is of clear ecological importance.

In summary, our findings show that the analysis of natural communities using network analysis represents an ideal way of visualizing the complexity present within these communities, but also represents a manner of mechanistically representing the differences observed across the reproductive success of individuals and/or species while linking them to potential ecological mechanisms. Our findings represent a step forward in our understanding of how community structure affects function, yet they also show that more studies with better resolved communities are needed, with a special focus being placed in evaluating reproductive success of a larger array of plant species.

## Supplementary material

https://www.biorxiv.org/content/10.1101/629931v6.supplementary-material

## Data accessibility

All the data used are available at: https://zenodo.org/account/settings/github/repository/ibartomeus/BeeFunData and the code used to generate all results can be found at: https://doi.org/10.5281/zenodo.3364037

## Acknowledgements

The authors would like to thank Oscar Aguado for identifying pollinator species. We would also like to acknowledge the efforts of the editor Cédric Gaucherel, as well as those of Michael Lattorff, Nicolas Deguines, Alberto Pascual, Nico Blüthgen, J. M. Bennett and six anonymous reviewers who provided very helpful comments and suggestions in a previous version of this manuscript. AM received funding from a Juan de la Cierva (IJCI-2014-22558) and Ikerbasque fellowships. IB acknowledges funding from MSC-PCIG14-GA-2013-631653 BeeFun Project. We thank Doñana’s Singular Scientific-Technical Infrastructure (ICTS-RBD) for access to the park. Research also supported by the Spanish State Research Agency through María de Maeztu Excellence Unit accreditation (MDM-2017-0714) and the Basque Government BERC Programme.

Version 5 of this preprint has been peer-reviewed and recommended by Peer Community In Ecology (10.1101/629931).

## Conflict of interest disclosure

The authors of this preprint declare that they have no financial conflict of interest with the content of this article. AM and IB are recommenders for PCI Ecology.

## Statement of authorship

AM led data analysis, FPM collected data, IB obtained funding and supervised data collection and analysis. All authors contributed to writing

## Notes

https://zenodo.org/account/settings/github/repository/ibartomeus/BeeFunData

https://doi.org/10.5281/zenodo.3364037

### Competing Interest Statement

The authors have declared no competing interest.

### Summary of Updates

-include conflict of interest disclosure -link to PCI Ecology recommendation

