## Supplementary files and tables for "Niche complementarity among pollinators increases community-level plant reproductive success"

Short running title: Network structure effects on ecosystem functioning

^1^Basque Centre for Climate Change-BC3, Edif. Sede 1, 1º, Parque Científico UPV-EHU, Barrio Sarriena s/n, 48940, Leioa, Spain

^2^IKERBASQUE, Basque Foundation for Science, María Díaz de Haro 3, 48013, Bilbao, Spain

^3^Estación Biológica de Doñana (EBD-CSIC), Avda. Américo Vespucio 26, Isla de la Cartuja, 41092, Sevilla, Spain

### Supplementary material

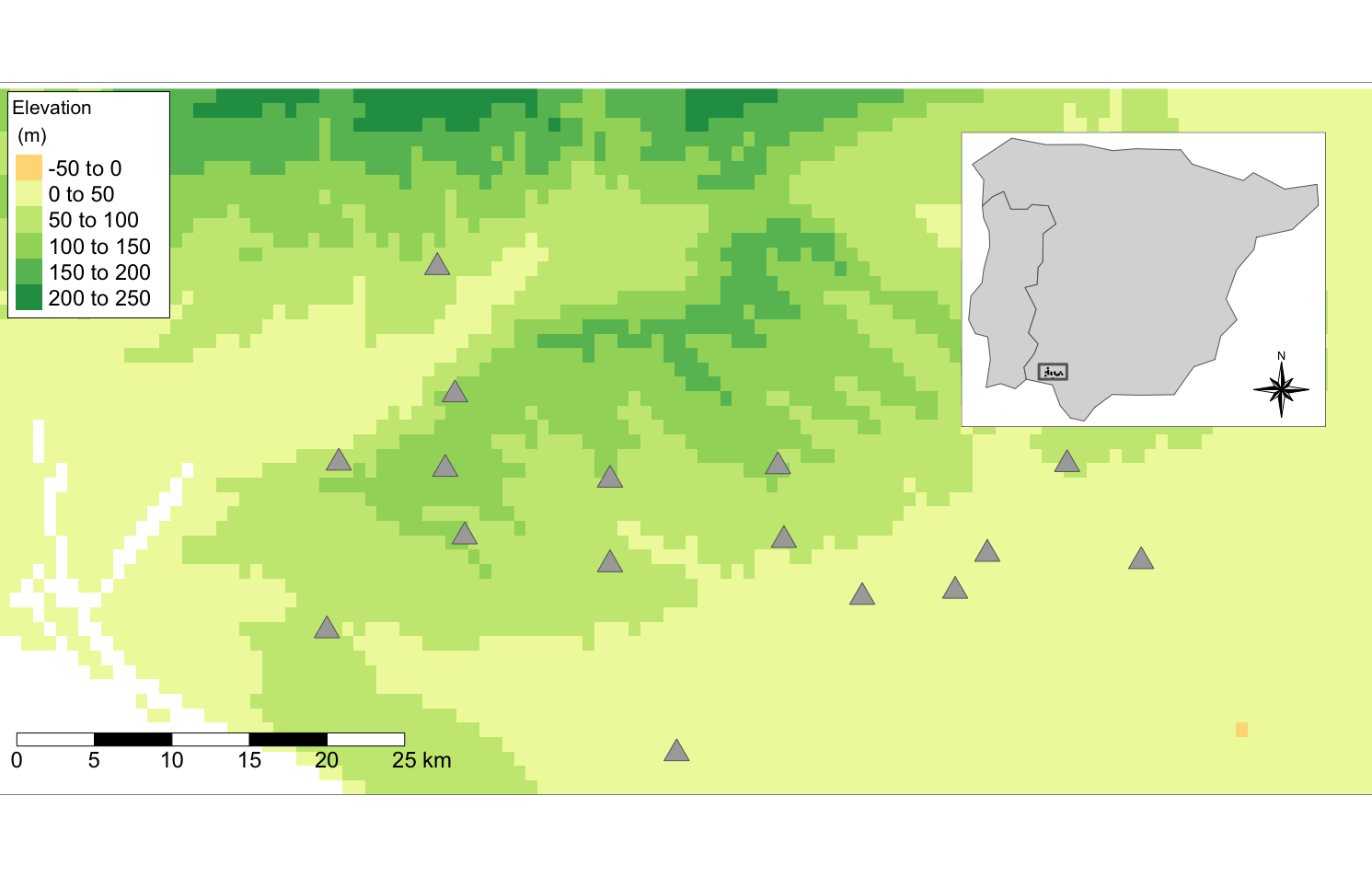

**Figure S1.** Map showing location of 16 Mediterranean woodland patches where plant-pollinator interactions were surveyed from February to May 2015. Inset shows location of study area within SW Spain.

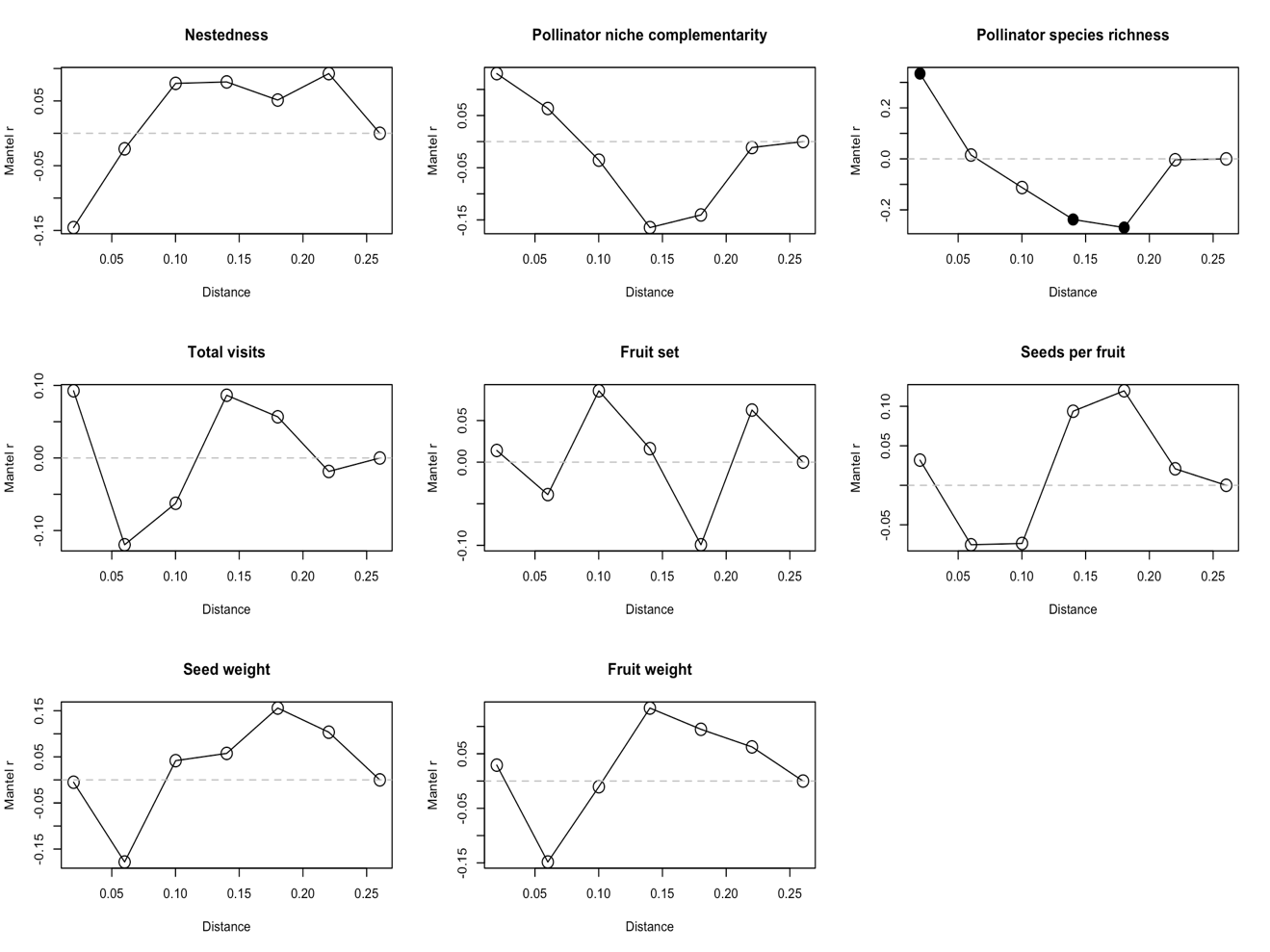

**Figure S2**. Mantel correlograms showing a low spatial autocorrelation for the different variables included in the analyses.

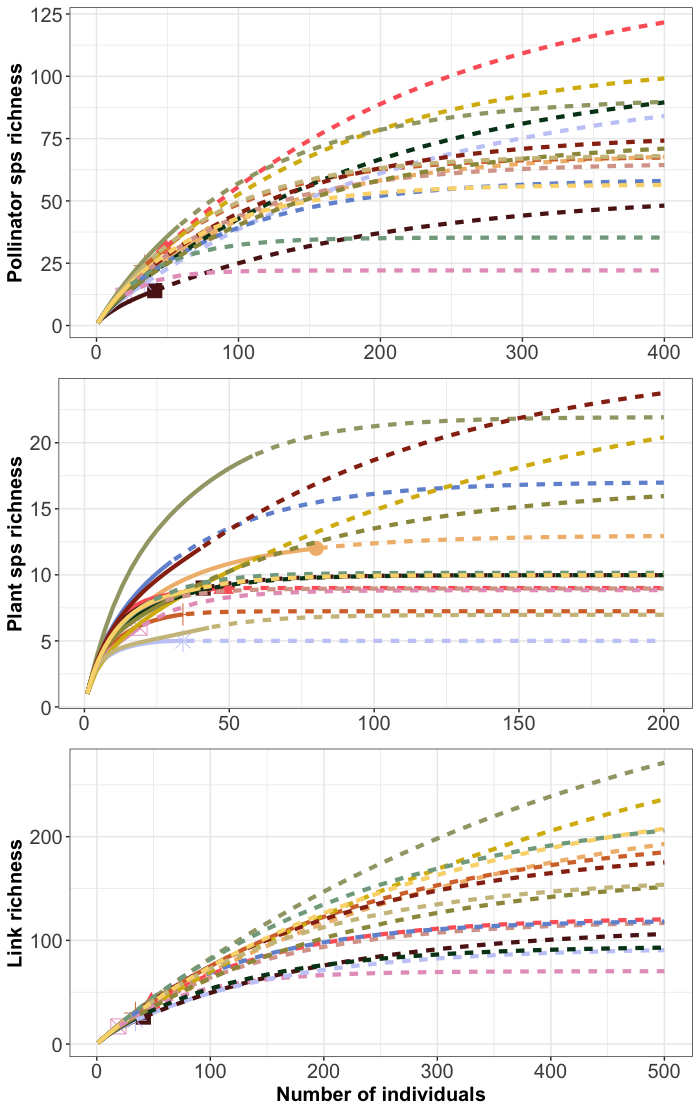

**Figure S3.** Accumulation curves of pollinator, plant and plant-pollinator link richness with increasing sampling effort up to 100% sample coverage. Solid lines and points indicate observed richness while dashed lines show expected richness at increasing sample size, (i.e., extrapolated).

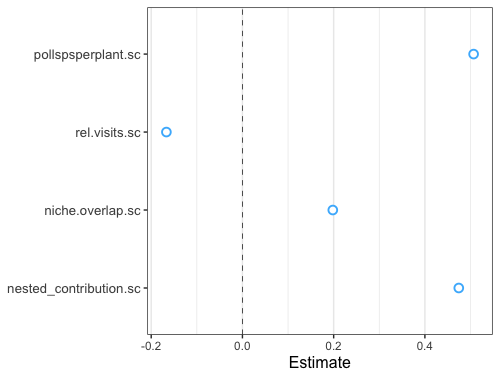

**Figure S4.** Forest plots showing 95% confidence intervals for all coefficients included in species-level models.

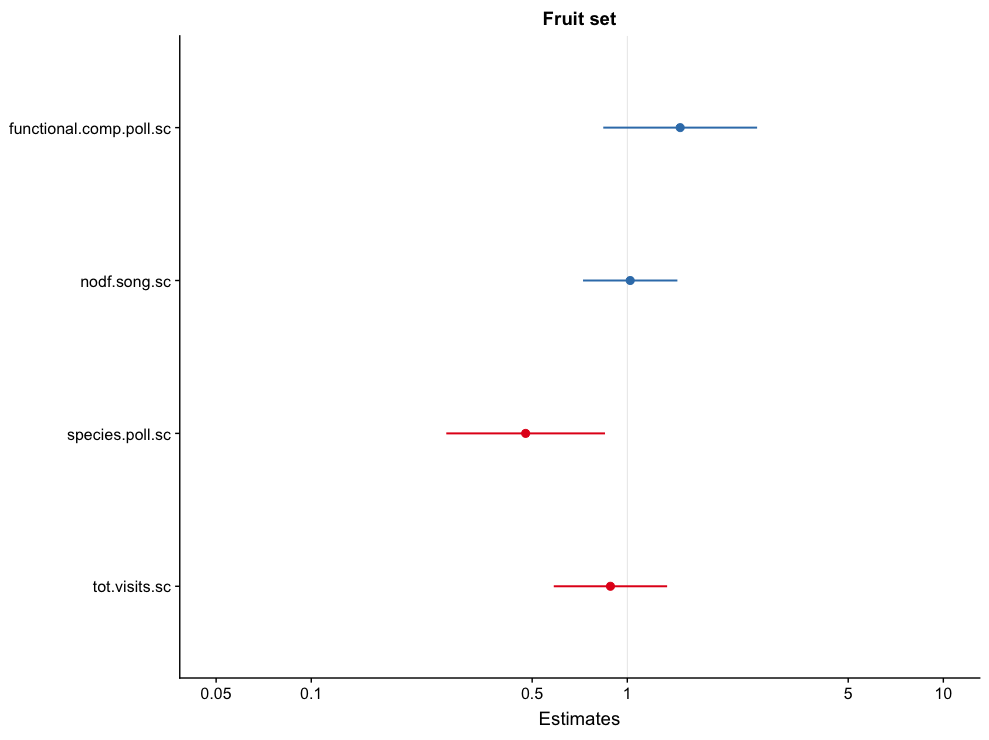

A)

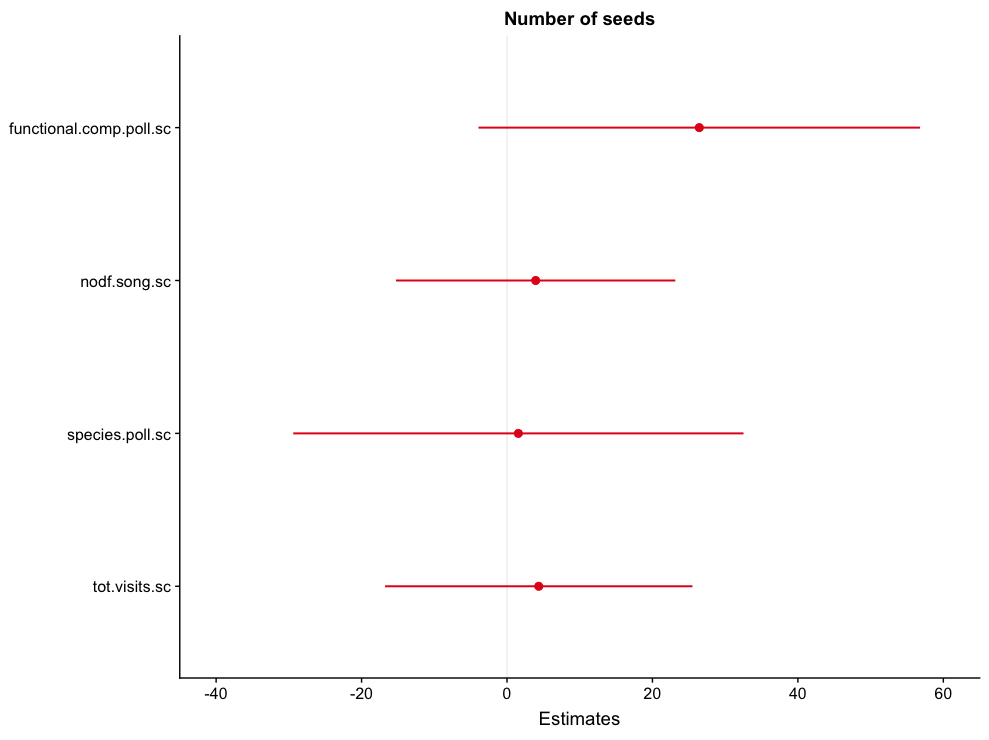

B)

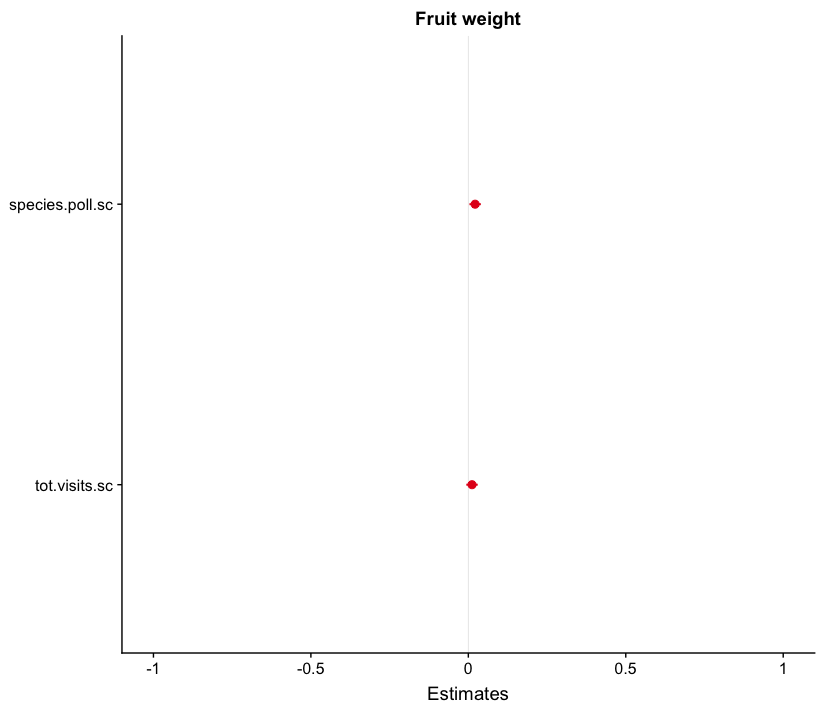

C)

**Figure S5**. Forest plots showing 95% confidence intervals for all coefficients included in site-level models, A) fruit set, B) number of seeds per fruit and C) fruit and seed weight.

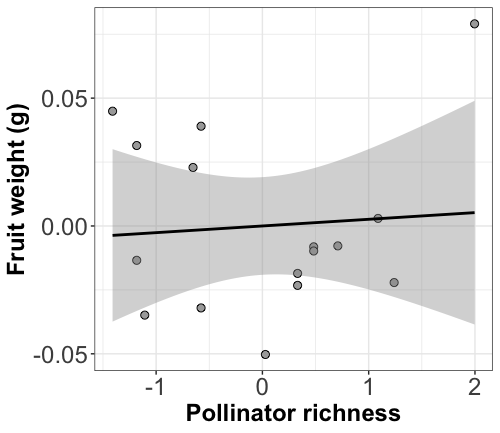

**Figure S6.** Partial residual plots showing the effect of pollinator richness on site-level average fruit weight. Dots represent values for each site (N=16 sites).

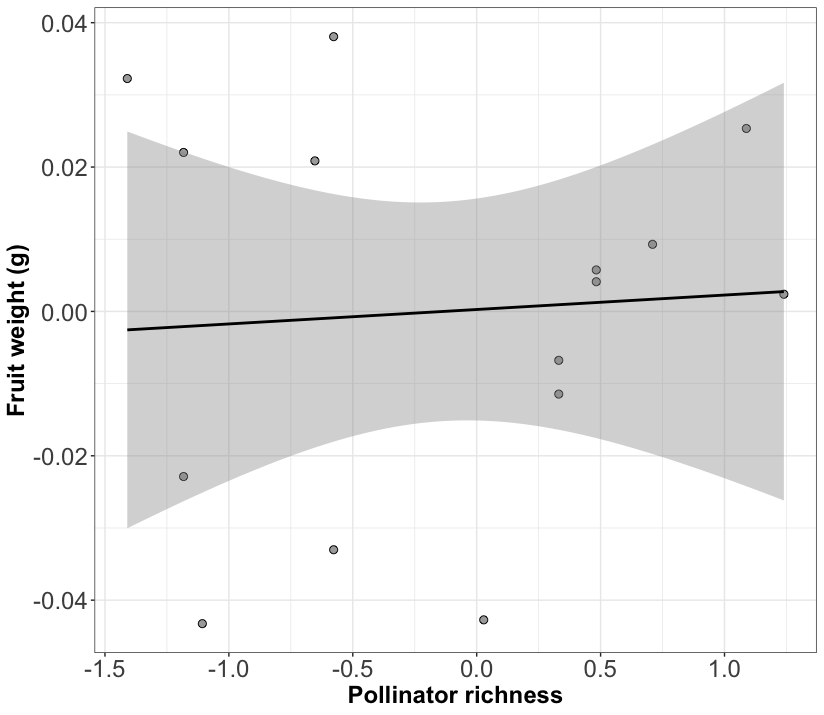

**Figure S7.** Partial residual plots showing the effect of pollinator richness on site-level average fruit weight. Here, a site with a particularly large pollinator richness value is removed to test whether it might be driving the significant relationship. Dots represent values for each site (N=15 sites).

**Table S1A.** List of all plant species present at each of the sites and included in network analyses.

| **Site** | **Plant species** |
| --- | --- |
| Aznalcazar | *Asphodelus fistulosus* |
| Aznalcazar | *Cistus crispus* |
| Aznalcazar | *Cistus ladanifer* |
| Aznalcazar | *Cistus monspeliensis* |
| Aznalcazar | *Cistus salvifolius* |
| Aznalcazar | *Echium plantagineum* |
| Aznalcazar | *Lavandula pedunculata* |
| Aznalcazar | *Lavandula stoechas* |
| Aznalcazar | *Lavatera cretica* |
| Aznalcazar | *Rosmarinus officinalis* |
| Aznalcazar | *Teucrium fruticans* |
| Bonares | *Andryala integrifolia* |
| Bonares | *Cistus crispus* |
| Bonares | *Cistus ladanifer* |
| Bonares | *Cistus salvifolius* |
| Bonares | *Halimium commutatum* |
| Bonares | *Lavandula pedunculata* |
| Bonares | *Lavandula stoechas* |
| Bonares | *Thapsia villosa* |
| Bonares | *Thymus mastichina* |
| ConventodelaLuz | *Cistus crispus* |
| ConventodelaLuz | *Cistus ladanifer* |
| ConventodelaLuz | *Cistus salvifolius* |
| ConventodelaLuz | *Halimium halimifolium* |
| ConventodelaLuz | *Lavandula stoechas* |
| ConventodelaLuz | *Retama* sp. |
| ConventodelaLuz | *Rosmarinus officinalis* |
| ConventodelaLuz | *Spartium junceum* |
| ConventodelaLuz | *Teucrium fruticans* |
| CotitodeSantaTeresa | *Astragalus lusitanicus* |
| CotitodeSantaTeresa | *Cistus crispus* |
| CotitodeSantaTeresa | *Cistus salvifolius* |
| CotitodeSantaTeresa | *Lavandula pedunculata* |
| CotitodeSantaTeresa | *Lavandula stoechas* |
| CotitodeSantaTeresa | *Rosmarinus officinalis* |
| CotitodeSantaTeresa | *Thapsia villosa* |
| Elpinar | *Cistus albidus* |
| Elpinar | *Cistus salvifolius* |
| Elpinar | *Convolvulus arvensis* |
| Elpinar | *Halimium commutatum* |
| Elpinar | *Lavandula stoechas* |
| Elpinar | *Rosmarinus officinalis* |
| Elpozo | *Cistus ladanifer* |
| Elpozo | *Cistus salvifolius* |
| Elpozo | *Erica scoparia* |
| Elpozo | *Erica umbellata* |
| Elpozo | *Rosmarinus officinalis* |
| Esparragal | *Armeria velutina* |
| Esparragal | *Chamaemelum fuscatum* |
| Esparragal | *Cistus libanotis* |
| Esparragal | *Cistus salvifolius* |
| Esparragal | *Halimium commutatum* |
| Esparragal | *Lavandula pedunculata* |
| Esparragal | *Lavandula stoechas* |
| Esparragal | *Scabiosa atropurpurea* |
| LaCunya | *Andryala integrifolia* |
| LaCunya | *Cerinthe gymnandra* |
| LaCunya | *Cistus salvifolius* |
| LaCunya | *Echium plantagineum* |
| LaCunya | *Erica ciliaris* |
| LaCunya | *Halimium commutatum* |
| LaCunya | *Lavandula pedunculata* |
| LaCunya | *Leontodon longirostris* |
| LaCunya | *Rosmarinus officinalis* |
| LaCunya | *Tuberaria guttata* |
| LaCunya | *Ulex australis* |
| LaRocina | *Anchusa azurea* |
| LaRocina | *Andryala integrifolia* |
| LaRocina | *Cistus salvifolius* |
| LaRocina | *Diplotaxis virgata* |
| LaRocina | *Halimium commutatum* |
| LaRocina | *Halimium halimifolium* |
| LaRocina | *Lavandula pedunculata* |
| LaRocina | *Lavandula stoechas* |
| LaRocina | *Linaria viscosa* |
| LaRocina | *Rosmarinus officinalis* |
| LaRocina | *Spartium junceum* |
| Lasmulas | *Cistus crispus* |
| Lasmulas | *Cistus ladanifer* |
| Lasmulas | *Cistus monspeliensis* |
| Lasmulas | *Cistus salvifolius* |
| Lasmulas | *Echium plantagineum* |
| Lasmulas | *Lavandula stoechas* |
| Lasmulas | *Ranunculus* sp. |
| Lasmulas | *Rosmarinus officinalis* |
| Lasmulas | *Thapsia villosa* |
| Niebla | *Andryala integrifolia* |
| Niebla | *Arctotheca calendula* |
| Niebla | *Asphodelus fistulosus* |
| Niebla | *Astragalus lusitanicus* |
| Niebla | *Calendula arvensis* |
| Niebla | *Carduus* sp. |
| Niebla | *Cistus crispus* |
| Niebla | *Cistus ladanifer* |
| Niebla | *Cistus monspeliensis* |
| Niebla | *Convolvulus arvensis* |
| Niebla | *Lavandula pedunculata* |
| Niebla | *Lavandula stoechas* |
| Niebla | *Leontodon* sp. |
| Niebla | *Linaria viscosa* |
| Niebla | *Linum bienne* |
| Niebla | *Lupinus angustifolius* |
| Niebla | *Phlomis purpurea* |
| Niebla | *Taraxacum vulgare* |
| Niebla | *Thapsia villosa* |
| PinaresdeHinojos | *Andryala integrifolia* |
| PinaresdeHinojos | *Cistus crispus* |
| PinaresdeHinojos | *Cistus libanotis* |
| PinaresdeHinojos | *Cistus salvifolius* |
| PinaresdeHinojos | *Diplotaxis virgata* |
| PinaresdeHinojos | *Rosmarinus officinalis* |
| PinaresdeHinojos | *Spartium junceum* |
| PinaresdeHinojos | *Ulex australis* |
| Pinodelcuervo | *Asphodelus fistulosus* |
| Pinodelcuervo | *Chamaemelum fuscatum* |
| Pinodelcuervo | *Cistus crispus* |
| Pinodelcuervo | *Cistus ladanifer* |
| Pinodelcuervo | *Cistus salvifolius* |
| Pinodelcuervo | *Halimium commutatum* |
| Pinodelcuervo | *Lavandula pedunculata* |
| Pinodelcuervo | *Lavandula stoechas* |
| Pinodelcuervo | *Ranunculus* sp. |
| Pinodelcuervo | *Rosmarinus officinalis* |
| Pinodelcuervo | *Thapsia villosa* |
| Pinodelcuervo | *Ulex australis* |
| Urbanizaciones | *Calendula arvensis* |
| Urbanizaciones | *Cistus crispus* |
| Urbanizaciones | *Cistus salvifolius* |
| Urbanizaciones | *Halimium commutatum* |
| Urbanizaciones | *Lavandula pedunculata* |
| Urbanizaciones | *Lavandula stoechas* |
| Urbanizaciones | *Rosmarinus officinalis* |
| Urbanizaciones | *Tuberaria guttata* |
| Urbanizaciones | *Ulex australis* |
| Villamanriqueeste | *Cistus crispus* |
| Villamanriqueeste | *Cistus ladanifer* |
| Villamanriqueeste | *Cistus salvifolius* |
| Villamanriqueeste | *Genista hirsuta* |
| Villamanriqueeste | *Rosmarinus officinalis* |
| Villamanriqueeste | *Spartium junceum* |
| Villamanriquesur | *Andryala integrifolia* |
| Villamanriquesur | *Armeria velutina* |
| Villamanriquesur | *Cistus crispus* |
| Villamanriquesur | *Cistus salvifolius* |
| Villamanriquesur | *Convolvulus arvensis* |
| Villamanriquesur | *Genista hirsuta* |
| Villamanriquesur | *Halimium halimifolium* |
| Villamanriquesur | *Lavandula stoechas* |
| Villamanriquesur | *Rosmarinus officinalis* |

**Table S1B.** List of all pollinator species present at each of the sites and included in network analyses.

| **Site** | ***Pollinator species*** |
| --- | --- |
| Aznalcazar | *Andrena flavipes* |
| Aznalcazar | *Andrena nigroaenaea* |
| Aznalcazar | *Andrena nitidiuscula* |
| Aznalcazar | *Andrena* sp. |
| Aznalcazar | *Andrena tenuistriata* |
| Aznalcazar | *Anthophora dispar* |
| Aznalcazar | *Anthophora* sp. |
| Aznalcazar | *Apis mellifera* |
| Aznalcazar | *Bombus terrestris* |
| Aznalcazar | *Calliphora* sp. |
| Aznalcazar | *Cerceris sabulosa* |
| Aznalcazar | *Dasypoda argentata* |
| Aznalcazar | *Dasypoda cingulata* |
| Aznalcazar | *Dasypoda crassicornis* |
| Aznalcazar | *Empis* morpho1 |
| Aznalcazar | *Eristalis arbustorum* |
| Aznalcazar | *Eucera alternans* |
| Aznalcazar | *Eucera codinai* |
| Aznalcazar | *Eucera collaris* |
| Aznalcazar | *Eucera elongatula* |
| Aznalcazar | *Eucera hispaliensis* |
| Aznalcazar | *Eucera* sp. |
| Aznalcazar | *Flavipanurgus venustus* |
| Aznalcazar | *Heliotaurus ruficollis* |
| Aznalcazar | *Hoplitis adunca* |
| Aznalcazar | *Lasioglossum* morpho1 |
| Aznalcazar | *Macroglossum stellatarum* |
| Aznalcazar | *Merodon* sp. |
| Aznalcazar | *Osmia leaiana* |
| Aznalcazar | *Panurgus calcaratus* |
| Aznalcazar | *Pseudoanthidium lituratum* |
| Aznalcazar | *Rhyncomyia cuprea* |
| Aznalcazar | *Syrphidae* sp. |
| Aznalcazar | *Tabanus* morpho1 |
| Aznalcazar | *Tabanus* morpho2 |
| Aznalcazar | *Volucella elegans* |
| Aznalcazar | *Xylocopa cantabrita* |
| Bonares | *Ammophila heydeni* |
| Bonares | *Ancistrocerus biphaleratus* |
| Bonares | *Andrena hispania* |
| Bonares | *Andrena nigroaenaea* |
| Bonares | *Andrena ovatula* |
| Bonares | *Andrena rhyssonota* |
| Bonares | *Andrena vulpecula* |
| Bonares | *Anthaxia* morpho1 |
| Bonares | *Anthidium septemspinosum* |
| Bonares | *Apis mellifera* |
| Bonares | *Bombus terrestris* |
| Bonares | *Bombylius* sp. |
| Bonares | *Ceratina cucurbitina* |
| Bonares | *Colletes acutus* |
| Bonares | *Colletes ligatus* |
| Bonares | *Dasypoda hirtipes* |
| Bonares | *Dasypogon* morpho1 |
| Bonares | *Empis* morpho1 |
| Bonares | *Empis* sp. |
| Bonares | *Eristalis* sp. |
| Bonares | *Eucera collaris* |
| Bonares | *Eucera elongatula* |
| Bonares | *Eucera* sp. |
| Bonares | *Graphosoma lineatum* |
| Bonares | *Halictus scabiosae* |
| Bonares | *Hoplitis papaveris* |
| Bonares | *Lasioglossum* sp. |
| Bonares | *Megachile* sp. |
| Bonares | *Musca* sp. |
| Bonares | *Platynochaetus setosus* |
| Bonares | *Trypoxylon* morpho1 |
| ConventodelaLuz | *Ammophila heydeni* |
| ConventodelaLuz | *Anthophora retusa* |
| ConventodelaLuz | *Apis mellifera* |
| ConventodelaLuz | *Bombus terrestris* |
| ConventodelaLuz | *Chasmatopterus villosulus* |
| ConventodelaLuz | *Eucera alternans* |
| ConventodelaLuz | *Exosoma lusitanicum* |
| ConventodelaLuz | *Ichneumonidae* morpho1 |
| ConventodelaLuz | *Oxythyrea funesta* |
| ConventodelaLuz | *Platynochaetus setosus* |
| ConventodelaLuz | *Syrphidae* sp. |
| ConventodelaLuz | *Tropinota squalida* |
| ConventodelaLuz | *Vespula germanica* |
| ConventodelaLuz | *Xylocopa cantabrita* |
| CotitodeSantaTeresa | *Andrena rhyssonota* |
| CotitodeSantaTeresa | *Anthophora aestivalis* |
| CotitodeSantaTeresa | *Anthophora hispanica* |
| CotitodeSantaTeresa | *Apis mellifera* |
| CotitodeSantaTeresa | *Bombus terrestris* |
| CotitodeSantaTeresa | *Dasypoda cingulata* |
| CotitodeSantaTeresa | *Dasypoda crassicornis* |
| CotitodeSantaTeresa | *Eucera chrysopyga* |
| CotitodeSantaTeresa | *Eucera codinai* |
| CotitodeSantaTeresa | *Eucera* sp. |
| CotitodeSantaTeresa | *Heriades crenulatus* |
| CotitodeSantaTeresa | *Lasioglossum albocinctum* |
| CotitodeSantaTeresa | *Lasioglossum malachurum* |
| CotitodeSantaTeresa | *Lasioglossum* sp. |
| CotitodeSantaTeresa | *Lestica clypeata* |
| CotitodeSantaTeresa | *Macroglossum stellatarum* |
| CotitodeSantaTeresa | *Merodon* sp. |
| CotitodeSantaTeresa | *Musca* morpho1 |
| CotitodeSantaTeresa | *Nemotelus* morpho1 |
| CotitodeSantaTeresa | *Nomada agrestis* |
| CotitodeSantaTeresa | *Nomada* sp. |
| CotitodeSantaTeresa | *Platynochaetus setosus* |
| CotitodeSantaTeresa | *Trypoxylon* morpho1 |
| CotitodeSantaTeresa | *Xylocopa cantabrita* |
| Elpinar | *Andrena ferrugineicrus* |
| Elpinar | *Andrena nigroaenaea* |
| Elpinar | *Apis mellifera* |
| Elpinar | *Bombus terrestris* |
| Elpinar | *Ceratina cucurbitina* |
| Elpinar | *Empis tessellata* |
| Elpinar | *Eucera alternans* |
| Elpinar | *Eucera* sp. |
| Elpinar | *Lestica clypeata* |
| Elpinar | *Platynochaetus setosus* |
| Elpinar | *Psilothrix viridicoerulea* |
| Elpinar | *Xylocopa cantabrita* |
| Elpozo | *Andrena hispania* |
| Elpozo | *Andrena* sp. |
| Elpozo | *Apis mellifera* |
| Elpozo | *Bombus terrestris* |
| Elpozo | *Bombylius* morpho1 |
| Elpozo | *Bombylius* sp. |
| Elpozo | *Bombylius torquatus* |
| Elpozo | *Colletes nigricans* |
| Elpozo | *Dasypoda crassicornis* |
| Elpozo | *Heliotaurus ruficollis* |
| Elpozo | *Lasioglossum bimaculatus* |
| Elpozo | *Lasioglossum imminutus* |
| Elpozo | *Merodon* sp. |
| Elpozo | *Musca* sp. |
| Elpozo | *Panurgus* sp. |
| Elpozo | *Psilothrix viridicoerulea* |
| Elpozo | *Xylocopa violacea* |
| Esparragal | *Andrena* sp. |
| Esparragal | *Anthophora atroalba* |
| Esparragal | *Apidae* sp. |
| Esparragal | *Apis mellifera* |
| Esparragal | *Cerceris* morpho1 |
| Esparragal | *Chasmatopterus illigeri* |
| Esparragal | *Dasypoda* sp. |
| Esparragal | *Episyrphus balteatus* |
| Esparragal | *Eucera collaris* |
| Esparragal | *Halictus tridivisus* |
| Esparragal | *Lasioglossum bimaculatus* |
| Esparragal | *Lasioglossum leucozonium* |
| Esparragal | *Lasioglossum malachurum* |
| Esparragal | *Lasioglossum* morpho1 |
| Esparragal | *Osmia fulviventris* |
| Esparragal | *Pieris rapae* |
| Esparragal | *Tenthredo* sp. |
| Esparragal | *Usia* morpho1 |
| LaCunya | *Andrena rhyssonota* |
| LaCunya | *Anthophora dispar* |
| LaCunya | *Anthophora retusa* |
| LaCunya | *Apis mellifera* |
| LaCunya | *Bombus terrestris* |
| LaCunya | *Ceratina cucurbitina* |
| LaCunya | *Dasypoda cingulata* |
| LaCunya | *Empis* morpho1 |
| LaCunya | *Heliotaurus ruficollis* |
| LaCunya | *Lasioglossum albocinctum* |
| LaCunya | *Lasioglossum imminutus* |
| LaCunya | *Lasioglossum malachurum* |
| LaCunya | *Lasioglossum* sp. |
| LaCunya | *Lasioglossum tridivisus* |
| LaCunya | *Lomatia* morpho1 |
| LaCunya | *Panurgus banksianus* |
| LaCunya | *Pieris brassicae* |
| LaCunya | *Pseudoanthidium melanorum* |
| LaRocina | *Andrena* sp. |
| LaRocina | *Anthophora bimaculata* |
| LaRocina | *Anthophora retusa* |
| LaRocina | *Apis mellifera* |
| LaRocina | *Arachnospila* morpho1 |
| LaRocina | *Bombus terrestris* |
| LaRocina | *Ceratina* sp. |
| LaRocina | *Colletes acutus* |
| LaRocina | *Colletes* sp. |
| LaRocina | *Dasypoda cingulata* |
| LaRocina | *Dasypoda crassicornis* |
| LaRocina | *Dasypoda* sp. |
| LaRocina | *Dischistus morpho1* |
| LaRocina | *Dischistus senex* |
| LaRocina | *Episyrphus balteatus* |
| LaRocina | *Eristalis tenax* |
| LaRocina | *Helophilus trivittatus* |
| LaRocina | *Heriades crenulatus* |
| LaRocina | *Heriades truncorum* |
| LaRocina | *Hoplitis tridentata* |
| LaRocina | *Lasioglossum imminutus* |
| LaRocina | *Lasioglossum malachurum* |
| LaRocina | *Lasioglossum* sp. |
| LaRocina | *Malachius* morpho1 |
| LaRocina | *Merodon* sp. |
| LaRocina | *Nomada agrestis* |
| LaRocina | *Nomada fucata* |
| LaRocina | *Nomada melathoracica* |
| LaRocina | *Osmia caerulescens* |
| LaRocina | *Panurgus banksianus* |
| LaRocina | *Panurgus* sp. |
| LaRocina | *Rhyncomyia cuprea* |
| LaRocina | *Sphecodes* sp. |
| LaRocina | *Syrphidae* sp. |
| LaRocina | *Xylocopa cantabrita* |
| Lasmulas | *Andrena flavipes* |
| Lasmulas | *Andrena nigroaenaea* |
| Lasmulas | *Andrena rhyssonota* |
| Lasmulas | *Anthophora dispar* |
| Lasmulas | *Anthophora hispanica* |
| Lasmulas | *Apis mellifera* |
| Lasmulas | *Bombylius* sp. |
| Lasmulas | *Bombylius torquatus* |
| Lasmulas | *Conopidae* sp. |
| Lasmulas | *Dasypoda albimana* |
| Lasmulas | *Empis* sp. |
| Lasmulas | *Empis testacea* |
| Lasmulas | *Eristalis similis* |
| Lasmulas | *Eucera chrysopyga* |
| Lasmulas | *Eucera* sp. |
| Lasmulas | *Flavipanurgus venustus* |
| Lasmulas | *Heliotaurus ruficollis* |
| Lasmulas | *Lasioglossum imminutus* |
| Lasmulas | *Lasioglossum malachurum* |
| Lasmulas | *Mycterus curculioides* |
| Lasmulas | *Panurgus calcaratus* |
| Lasmulas | *Panurgus dargius* |
| Lasmulas | *Xylocopa cantabrita* |
| Niebla | *Andrena flavipes* |
| Niebla | *Andrena labialis* |
| Niebla | *Andrena ovatula* |
| Niebla | *Andrena rhyssonota* |
| Niebla | *Andrena tenuistriata* |
| Niebla | *Anthidium septemspinosum* |
| Niebla | *Anthophora dispar* |
| Niebla | *Anthophora hispanica* |
| Niebla | *Anthophora* sp. |
| Niebla | *Apis mellifera* |
| Niebla | *Bombus terrestris* |
| Niebla | *Bombylius fimbriatus* |
| Niebla | *Bombylius* sp. |
| Niebla | *Ceratina callosa* |
| Niebla | *Colletes* sp. |
| Niebla | *Episyrphus balteatus* |
| Niebla | *Eucera collaris* |
| Niebla | *Eucera notata* |
| Niebla | *Exosoma lusitanicum* |
| Niebla | *Halictus scabiosae* |
| Niebla | *Heliotaurus ruficollis* |
| Niebla | *Heriades crenulatus* |
| Niebla | *Lasioglossum malachurum* |
| Niebla | *Lasioglossum* sp. |
| Niebla | *Macrophya montana* |
| Niebla | *Merodon* sp. |
| Niebla | *Osmia bicornis* |
| Niebla | *Osmia submicans* |
| Niebla | *Panurgus banksianus* |
| Niebla | *Panurgus dargius* |
| Niebla | *Platynochaetus setosus* |
| Niebla | *Potosia cuprea* |
| Niebla | *Rhodanthidium sticticum* |
| Niebla | *Sphaerophoria scripta* |
| Niebla | *Systropha planidens* |
| Niebla | *Usia* morpho1 |
| Niebla | *Usia* morpho2 |
| Niebla | *Usia* sp. |
| Niebla | *Vespula germanica* |
| Niebla | *Xylocopa violacea* |
| PinaresdeHinojos | *Andrena hispania* |
| PinaresdeHinojos | *Apis mellifera* |
| PinaresdeHinojos | *Bombus terrestris* |
| PinaresdeHinojos | *Chrysura refulgens* |
| PinaresdeHinojos | *Colletes acutus* |
| PinaresdeHinojos | *Colletes nigricans* |
| PinaresdeHinojos | *Colletes* sp. |
| PinaresdeHinojos | *Dasypoda crassicornis* |
| PinaresdeHinojos | *Lasioglossum bimaculatus* |
| PinaresdeHinojos | *Lasioglossum malachurum* |
| PinaresdeHinojos | *Lasioglossum* sp. |
| PinaresdeHinojos | *Nomada melathoracica* |
| PinaresdeHinojos | *Panurgus dargius* |
| PinaresdeHinojos | *Psilothrix viridicoerulea* |
| PinaresdeHinojos | *Tenthredo corynetes* |
| PinaresdeHinojos | *Xylocopa cantabrita* |
| Pinodelcuervo | *Ancistrocerus gazella* |
| Pinodelcuervo | *Ancistrocerus reconditus* |
| Pinodelcuervo | *Andrena hispania* |
| Pinodelcuervo | *Andrena* sp. |
| Pinodelcuervo | *Apis mellifera* |
| Pinodelcuervo | *Bombylella atra* |
| Pinodelcuervo | *Bombylius* sp. |
| Pinodelcuervo | *Ceratina cucurbitina* |
| Pinodelcuervo | *Cerceris* morpho1 |
| Pinodelcuervo | *Dasypoda cingulata* |
| Pinodelcuervo | *Flavipanurgus venustus* |
| Pinodelcuervo | *Lasioglossum sexnotatum* |
| Pinodelcuervo | *Lomatia* morpho1 |
| Pinodelcuervo | *Megascolia maculata* |
| Pinodelcuervo | *Merodon* sp. |
| Pinodelcuervo | *Musca* sp. |
| Pinodelcuervo | *Nomada melathoracica* |
| Pinodelcuervo | *Nomada merceti* |
| Pinodelcuervo | *Nomada* sp. |
| Pinodelcuervo | *Panurgus cephalotes* |
| Pinodelcuervo | *Pelecocera tricincta* |
| Pinodelcuervo | *Systoechus* morpho1 |
| Pinodelcuervo | *Usia* sp. |
| Pinodelcuervo | *Xylocopa cantabrita* |
| Urbanizaciones | *Andrena* sp. |
| Urbanizaciones | *Andrena vulpecula* |
| Urbanizaciones | *Apis mellifera* |
| Urbanizaciones | *Bombus terrestris* |
| Urbanizaciones | *Bombylidae* morpho1 |
| Urbanizaciones | *Bombylius* sp. |
| Urbanizaciones | *Ceratina* sp. |
| Urbanizaciones | *Colletes nigricans* |
| Urbanizaciones | *Dasypoda cingulata* |
| Urbanizaciones | *Dasypoda* sp. |
| Urbanizaciones | *Dischistus senex* |
| Urbanizaciones | *Eristalis tenax* |
| Urbanizaciones | *Eucera* sp. |
| Urbanizaciones | *Flavipanurgus venustus* |
| Urbanizaciones | *Lasioglossum albocinctum* |
| Urbanizaciones | *Lasioglossum imminutus* |
| Urbanizaciones | *Malachius* morpho1 |
| Urbanizaciones | *Osmia submicans* |
| Urbanizaciones | *Sphaerophoria scripta* |
| Urbanizaciones | *Xylocopa cantabrita* |
| Villamanriqueeste | *Andrena fertoni* |
| Villamanriqueeste | *Andrena flavipes* |
| Villamanriqueeste | *Andrena hispania* |
| Villamanriqueeste | *Anthophora dispar* |
| Villamanriqueeste | *Anthophora* sp. |
| Villamanriqueeste | *Apis mellifera* |
| Villamanriqueeste | *Bembix oculata* |
| Villamanriqueeste | *Bombylella atra* |
| Villamanriqueeste | *Dasypoda albimana* |
| Villamanriqueeste | *Dasypoda cingulata* |
| Villamanriqueeste | *Dasypoda crassicornis* |
| Villamanriqueeste | *Dischistus senex* |
| Villamanriqueeste | *Episyrphus balteatus* |
| Villamanriqueeste | *Eristalis* sp. |
| Villamanriqueeste | *Eucera collaris* |
| Villamanriqueeste | *Eumenes coarctatus* |
| Villamanriqueeste | *Flavipanurgus venustus* |
| Villamanriqueeste | *Helophilus* sp. |
| Villamanriqueeste | *Lasioglossum imminutus* |
| Villamanriqueeste | *Lasioglossum malachurum* |
| Villamanriqueeste | *Lasioglossum* sp. |
| Villamanriqueeste | *Musca* sp. |
| Villamanriqueeste | *Oxythyrea funesta* |
| Villamanriqueeste | *Panurgus banksianus* |
| Villamanriqueeste | *Panurgus cephalotes* |
| Villamanriqueeste | *Panurgus dargius* |
| Villamanriqueeste | *Psilothrix viridicoerulea* |
| Villamanriqueeste | *Vespula germanica* |
| Villamanriqueeste | *Xylocopa cantabrita* |
| Villamanriquesur | *Andrena flavipes* |
| Villamanriquesur | *Andrena nigroaenaea* |
| Villamanriquesur | *Apis mellifera* |
| Villamanriquesur | *Bibio* sp. |
| Villamanriquesur | *Calliphora* sp. |
| Villamanriquesur | *Dasypoda albimana* |
| Villamanriquesur | *Dasypoda cingulata* |
| Villamanriquesur | *Dasypoda crassicornis* |
| Villamanriquesur | *Dasypoda iberica* |
| Villamanriquesur | *Empis* sp. |
| Villamanriquesur | *Eristalinus taeniops* |
| Villamanriquesur | *Eucera bolivari* |
| Villamanriquesur | *Eucera collaris* |
| Villamanriquesur | *Eucera* sp. |
| Villamanriquesur | *Exosoma lusitanicum* |
| Villamanriquesur | *Lasioglossum malachurum* |
| Villamanriquesur | *Machimus* sp. |
| Villamanriquesur | *Merodon* sp. |
| Villamanriquesur | *Nomada* sp. |
| Villamanriquesur | *Pangonius micans* |
| Villamanriquesur | *Panurgus banksianus* |
| Villamanriquesur | *Panurgus calcaratus* |
| Villamanriquesur | *Panurgus dargius* |
| Villamanriquesur | *Pieris rapae* |
| Villamanriquesur | *Sphaerophoria scripta* |

**Table S2.** Number of individuals per plant species sampled at each site to assess reproductive success.

| Site | Plant species | Number of individuals |
| --- | --- | --- |
| LaRocina | *Anchusa azurea* | 6 |
| Aznalcazar | *Asphodelus fistulosus* | 9 |
| Niebla | *Asphodelus fistulosus* | 9 |
| Pinodelcuervo | *Asphodelus fistulosus* | 6 |
| CotitodeSantaTeresa | *Astragalus lusitanicus* | 2 |
| CotitodeSantaTeresa | *Cistus albidus* | 6 |
| Bonares | *Cistus crispus* | 6 |
| Niebla | *Cistus crispus* | 5 |
| Pinodelcuervo | *Cistus crispus* | 6 |
| Villamanriquesur | *Cistus crispus* | 3 |
| Aznalcazar | *Cistus ladanifer* | 6 |
| Bonares | *Cistus ladanifer* | 12 |
| ConventodelaLuz | *Cistus ladanifer* | 9 |
| Lasmulas | *Cistus ladanifer* | 6 |
| Niebla | *Cistus ladanifer* | 9 |
| Pinodelcuervo | *Cistus ladanifer* | 9 |
| Villamanriquesur | *Cistus ladanifer* | 7 |
| Lasmulas | *Cistus libanotis* | 6 |
| PinaresdeHinojos | *Cistus libanotis* | 6 |
| Niebla | *Cistus monspeliensis* | 3 |
| Aznalcazar | *Cistus salvifolius* | 9 |
| Bonares | *Cistus salvifolius* | 9 |
| ConventodelaLuz | *Cistus salvifolius* | 6 |
| CotitodeSantaTeresa | *Cistus salvifolius* | 5 |
| Esparragal | *Cistus salvifolius* | 3 |
| LaCunya | *Cistus salvifolius* | 6 |
| Lasmulas | *Cistus salvifolius* | 7 |
| PinaresdeHinojos | *Cistus salvifolius* | 6 |
| Urbanizaciones | *Cistus salvifolius* | 6 |
| Villamanriqueeste | *Cistus salvifolius* | 3 |
| Bonares | *Halimium commutatum* | 6 |
| Esparragal | *Halimium commutatum* | 7 |
| LaRocina | *Halimium commutatum* | 3 |
| Lasmulas | *Halimium commutatum* | 2 |
| Pinodelcuervo | *Halimium commutatum* | 6 |
| Urbanizaciones | *Halimium commutatum* | 6 |
| ConventodelaLuz | *Halimium halimifolium* | 3 |
| Esparragal | *Halimium halimifolium* | 3 |
| LaRocina | *Halimium halimifolium* | 3 |
| Villamanriquesur | *Halimium halimifolium* | 6 |
| Aznalcazar | *Lavandula pedunculata* | 4 |
| Esparragal | *Lavandula pedunculata* | 6 |
| LaCunya | *Lavandula pedunculata* | 5 |
| Niebla | *Lavandula pedunculata* | 9 |
| Bonares | *Lavandula stoechas* | 9 |
| ConventodelaLuz | *Lavandula stoechas* | 3 |
| CotitodeSantaTeresa | *Lavandula stoechas* | 3 |
| Lasmulas | *Lavandula stoechas* | 9 |
| Urbanizaciones | *Lavandula stoechas* | 9 |
| Villamanriquesur | *Lavandula stoechas* | 3 |
| Niebla | *Phlomis purpurea* | 6 |
| PinaresdeHinojos | *Retama sphaerocarpa* | 1 |
| ConventodelaLuz | *Rosmarinus officinalis* | 9 |
| CotitodeSantaTeresa | *Rosmarinus officinalis* | 6 |
| Elpinar | *Rosmarinus officinalis* | 8 |
| Elpozo | *Rosmarinus officinalis* | 6 |
| LaCunya | *Rosmarinus officinalis* | 6 |
| LaRocina | *Rosmarinus officinalis* | 1 |
| Pinodelcuervo | *Rosmarinus officinalis* | 9 |
| Urbanizaciones | *Rosmarinus officinalis* | 9 |
| Elpozo | *Spartium junceum* | 6 |
| LaRocina | *Spartium junceum* | 3 |
| ConventodelaLuz | *Teucrium fruticans* | 9 |
| Bonares | *Ulex australis* | 6 |
| Pinodelcuervo | *Ulex australis* | 3 |

**Table S3.** Number of fruits per plant species sampled at each site.

| Site | Plant species | Number of fruits |
| --- | --- | --- |
| Aznalcazar | *Asphodelus fistulosus* | 22 |
| Aznalcazar | *Cistus ladanifer* | 4 |
| Aznalcazar | *Cistus salvifolius* | 10 |
| Aznalcazar | *Lavandula pedunculata* | 10 |
| Bonares | *Cistus crispus* | 15 |
| Bonares | *Cistus ladanifer* | 12 |
| Bonares | *Cistus salvifolius* | 14 |
| Bonares | *Lavandula stoechas* | 23 |
| Bonares | *Ulex australis* | 21 |
| ConventodelaLuz | *Cistus ladanifer* | 4 |
| ConventodelaLuz | *Cistus salvifolius* | 5 |
| ConventodelaLuz | *Halimium halimifolium* | 6 |
| ConventodelaLuz | *Lavandula stoechas* | 13 |
| ConventodelaLuz | *Rosmarinus officinalis* | 23 |
| ConventodelaLuz | *Teucrium fruticans* | 16 |
| CotitodeSantaTeresa | *Astragalus lusitanicus* | 2 |
| CotitodeSantaTeresa | *Cistus albidus* | 8 |
| CotitodeSantaTeresa | *Cistus salvifolius* | 11 |
| CotitodeSantaTeresa | *Lavandula stoechas* | 11 |
| CotitodeSantaTeresa | *Rosmarinus officinalis* | 16 |
| Elpinar | *Rosmarinus officinalis* | 23 |
| Elpozo | *Rosmarinus officinalis* | 36 |
| Elpozo | *Spartium junceum* | 13 |
| Esparragal | *Cistus salvifolius* | 4 |
| Esparragal | *Halimium halimifolium* | 7 |
| Esparragal | *Lavandula pedunculata* | 16 |
| LaCunya | *Cistus salvifolius* | 8 |
| LaCunya | *Lavandula pedunculata* | 11 |
| LaCunya | *Rosmarinus officinalis* | 18 |
| LaRocina | *Anchusa azurea* | 10 |
| LaRocina | *Halimium commutatum* | 6 |
| LaRocina | *Halimium halimifolium* | 1 |
| LaRocina | *Rosmarinus officinalis* | 8 |
| LaRocina | *Spartium junceum* | 10 |
| Lasmulas | *Cistus ladanifer* | 5 |
| Lasmulas | *Cistus libanotis* | 10 |
| Lasmulas | *Cistus salvifolius* | 7 |
| Lasmulas | *Halimium commutatum* | 3 |
| Lasmulas | *Lavandula stoechas* | 23 |
| Niebla | *Asphodelus fistulosus* | 19 |
| Niebla | *Cistus crispus* | 10 |
| Niebla | *Cistus ladanifer* | 2 |
| Niebla | *Cistus monspeliensis* | 4 |
| Niebla | *Lavandula pedunculata* | 15 |
| Niebla | *Phlomis purpurea* | 23 |
| PinaresdeHinojos | *Cistus libanotis* | 9 |
| PinaresdeHinojos | *Cistus salvifolius* | 9 |
| PinaresdeHinojos | *Retama sphaerocarpa* | 2 |
| Pinodelcuervo | *Asphodelus fistulosus* | 10 |
| Pinodelcuervo | *Cistus crispus* | 18 |
| Pinodelcuervo | *Cistus ladanifer* | 10 |
| Pinodelcuervo | *Halimium commutatum* | 7 |
| Pinodelcuervo | *Rosmarinus officinalis* | 11 |
| Pinodelcuervo | *Ulex australis* | 5 |
| Urbanizaciones | *Cistus salvifolius* | 10 |
| Urbanizaciones | *Halimium commutatum* | 8 |
| Urbanizaciones | *Lavandula pedunculata* | 8 |
| Urbanizaciones | *Lavandula stoechas* | 16 |
| Urbanizaciones | *Rosmarinus officinalis* | 22 |
| Villamanriqueeste | *Cistus salvifolius* | 6 |
| Villamanriquesur | *Cistus crispus* | 11 |
| Villamanriquesur | *Cistus ladanifer* | 4 |
| Villamanriquesur | *Cistus salvifolius* | 4 |
| Villamanriquesur | *Halimium halimifolium* | 10 |
| Villamanriquesur | *Lavandula stoechas* | 8 |

**Table S4.** List of plant species surveyed and their mating system.

| **Plant_family** | **Plant_genus** | **Plant_species** | **reproductive_system** |
| --- | --- | --- | --- |
| Cistaceae | *Cistus* | *monspeliensis* | self-incompatible |
| Cistaceae | *Cistus* | *crispus* | self-incompatible |
| Cistaceae | *Cistus* | *ladanifer* | self-incompatible |
| Cistaceae | *Cistus* | *salviifolius* | self-incompatible |
| Cistaceae | *Cistus* | *albidus* | self-incompatible |
| Cistaceae | *Cistus* | *libanotis* | self-incompatible |
| Cistaceae | *Halimium* | *commutatum* | self-incompatible |
| Cistaceae | *Halimium* | *halimifolium* | self-incompatible |
| Lamiaceae | *Lavandula* | *pedunculata* | partially self-compatible |
| Lamiaceae | *Lavandula* | *stoechas* | partially self-compatible |
| Lamiaceae | *Teucrium* | *fruticans* | partially self-compatible |
| Lamiaceae | *Rosmarinus* | *officinalis* | partially self-compatible |
| Lamiaceae | *Phlomis* | *purpurea* | self-incompatible |
| Xanthorrhoeaceae | *Asphodelus* | *fistulosus* | partially self-compatible |
| Fabaceae | *Ulex* | *australis* | self-incompatible |
| Fabaceae | *Spartium* | *junceum* | self-incompatible |
| Fabaceae | *Astragalus* | *lusitanicus* | partially self-compatible |
| Fabaceae | *Retama* | *sphaerocarpa* | partially self-compatible |
| Boraginaceae | *Anchusa* | *azurea* | self-incompatible |

**Table S5.** Results of GLM showing effect of simple visitation metrics on A) site-level average fruit weight based on best model selected and B) the same analysis removing one site that has a particularly large pollinator richness value to test whether this point might be driving the relationship.

| A) | Estimate | Std. Error | t value |
| --- | --- | --- | --- |
| (Intercept) | 0.08 | 0.01 | 8.56 |
| **Pollinator richness** | **0.02** | **0.01** | **2.11** |
| Relative number of visits | 0.01 | 0.01 | 0.78 |

| B) | Estimate | Std. Error | t value |
| --- | --- | --- | --- |
| (Intercept) | 0.08 | 0.01 | 8.04 |
| **Pollinator richness** | **0.02** | **0.01** | **1.97** |
| Relative number of visits | 0.01 | 0.01 | 0.76 |

**Table S6.** Results of GLM showing effect of simple visitation metrics on equity in reproductive success across plant species within a site based on best model selected (0.50 threshold).

|  | Estimate | Std. Error | z value |
| --- | --- | --- | --- |
| (Intercept) | 0.40 | 0.63 | 0.64 |
| Pollinator richness | -0.41 | 1.18 | -0.35 |
| Relative number of visits | -0.28 | 0.72 | -0.39 |
| Nestedness | 0.87 | 1.02 | 0.86 |
| Pollinator niche complementarity | -0.51 | 1.32 | -0.38 |
